# Secure data storage on DNA hard drives

**DOI:** 10.1101/857748

**Authors:** Kaikai Chen, Jinbo Zhu, Filip Boskovic, Ulrich F. Keyser

## Abstract

DNA is emerging as a novel material for digital data storage. The two main challenges are efficient encoding and data security. Here, we develop an approach that allows for writing and erasing data by relying solely on Watson-Crick base pairing of short oligonucleotides to single-stranded DNA overhangs located along a long double-stranded DNA hard drive (DNA-HD). Our enzyme-free system enables fast synthesis-free data writing with predetermined building blocks. The use of DNA base pairing allows for secure encryption on DNA-HDs that requires a physical key and nanopore sensing for decoding. The system is suitable for miniature integration for an end-to-end DNA storage device. Our study opens a novel pathway for rewritable and secure data storage with DNA.

**One Sentence Summary:** Storing digital information on molecules along DNA hard drives for rewritable and secure data storage.

---

The importance of digital data is self-evident. Global digital data is growing exponentially with a never-ending trend, leading to demand beyond our ability to build media to store them (*1*). Currently, digital data is mainly stored in hard disk drives (HDDs), solid-state drives, optical disks and magnetic tapes. A large amount of energy is consumed to maintain and restore data because for most of these media the lifetime does not exceed 10 years. Alternatives are urgently needed for long-term and secure storage of important archival data.

Molecular data storage using DNA has become an emerging technology by encoding and writing data into the DNA sequence by synthesis (*2*). Data encoded in base pairs can be efficiently read by DNA sequencing. Synthesis-based DNA storage has advantages like high density, high durability and low maintenance costs (*3, 4*). Clelland et al. showed that DNA could be used to hide information as microdots (steganography) where they sent a message encoded on DNA (*5*). Church et al. encoded megabits data on DNA in 2012 showing the potential of DNA as the next-generation information storage material with a high data density (*6*). Around the same time, Goldman also stored and retrieved over five million bits of data on DNA (*7*). In recent years, DNA storage started booming with the amount of data stored in DNA raised from a few megabytes to hundreds of megabytes (*8*). However, fundamental challenges, including the complexity of DNA synthesis, still exist and have not been resolved even after years of optimization to achieve scalable and inexpensive data storage (*9*). Storing more data requires synthesis of ever more unique DNA strands, and it is unclear if this process can be significantly improved in the near future. In addition, storing data in the DNA sequence reaches the highest density but data is permanent and cannot be changed easily.

Therefore, alternatives are desperately needed which are not only for improving the encoding and writing strategy for the simple static data storage in DNA sequence but also for dynamic data operation such as rewritable capability and computing towards a realistic DNA computer. Employing existing native DNA strands rather than always synthesizing new strand are new routes, for instance, the DNA punch card method and the catalog DNA method (*10, 11*).

Data storage in 3D structures of DNA (*12-14*) or protein is another option as this may offer easier routes for data writing only by mixing components to build molecules. The information encoded in 3D structures of these molecules can be conveniently read by single-molecule methods such as nanopore sensing (*15-19*). We proposed a method for storing data on DNA nanostructures along double-stranded DNA (dsDNA) by mixing and incubating oligonucleotides and scaffolds (*20*). This method has intrinsic advantages in writing cost as we only need 2*n* oligonucleotide components for building libraries containing 2^n^ (∼5.2×10^33^ for *n*=112) different molecules. Our approach reduces the challenge to storage in DNA sequences where synthesising millions or billions of unique oligonucleotides is necessary.

Here, we dramatically improve the utility of DNA carriers by creating rewritable and secure storage data by reprogramming the individual oligonucleotides, inspired by the working principles of HDD. We name our system DNA hard drive (DNA-HD), which is easily operated only by mixing molecules at room temperature. Using our DNA-HDs, we can write data by attaching molecules on dsDNA and erase by targeted removal, and complete writing-erasing-rewriting cycle based only on Watson-Crick base pairing and toehold mediated strand displacement (*21-23*). More importantly, our system enables data encryption by physical keys (specific molecules bound to target positions) (*24*). Only addition of the correct molecular sequence enables decoding of the data combined with nanopore sensing, allowing for ultra-secure information storage on DNA. Our route to DNA data storage surpasses current approaches with advantages in efficient rewritable capability and data security.

We show the design of our platform in Figure 1. The DNA-HD is made by annealing the 7228 nucleotides (nt) linearised M13mp18 single-stranded DNA (ssDNA) scaffold (*25*) and complementary 38 oligonucleotides (Fig. 1A, Table S1, and Materials and Methods in the Supplementary). Oligonucleotides at designed positions are replaced with DNA dumbbells (*19*) and ssDNA overhangs. The DNA-HD is negatively charged so it can be read out using nanopore sensing by applying a potential to drive individual DNA-HDs through the sensing volume, which translate the molecular structure information into ionic current signals (Fig. 1B). Only unfolded translocations were retained for further analysis and the selection method is introduced in a previous study. When measured with our 10-15 nm nanopores a group of DNA dumbbells is able to create observable signals but the ssDNA overhang is too small to be detected (*26*). We employ the dumbbells at designed positions as references (REFs) to indicate the start and end of the reading, measure translocation velocity and determine the directionality of the DNA-HD. High (‘1’) bits are created by adding biotinylated oligonucleotides that hybridise to the ssDNA overhangs. Monovalent streptavidin binds to the biotin and creates an observable signal at the site. Thus we can encode data as ‘0’ (missing signal) and ‘1’ as peak is observed between the REF signals.

**Fig. 1.**
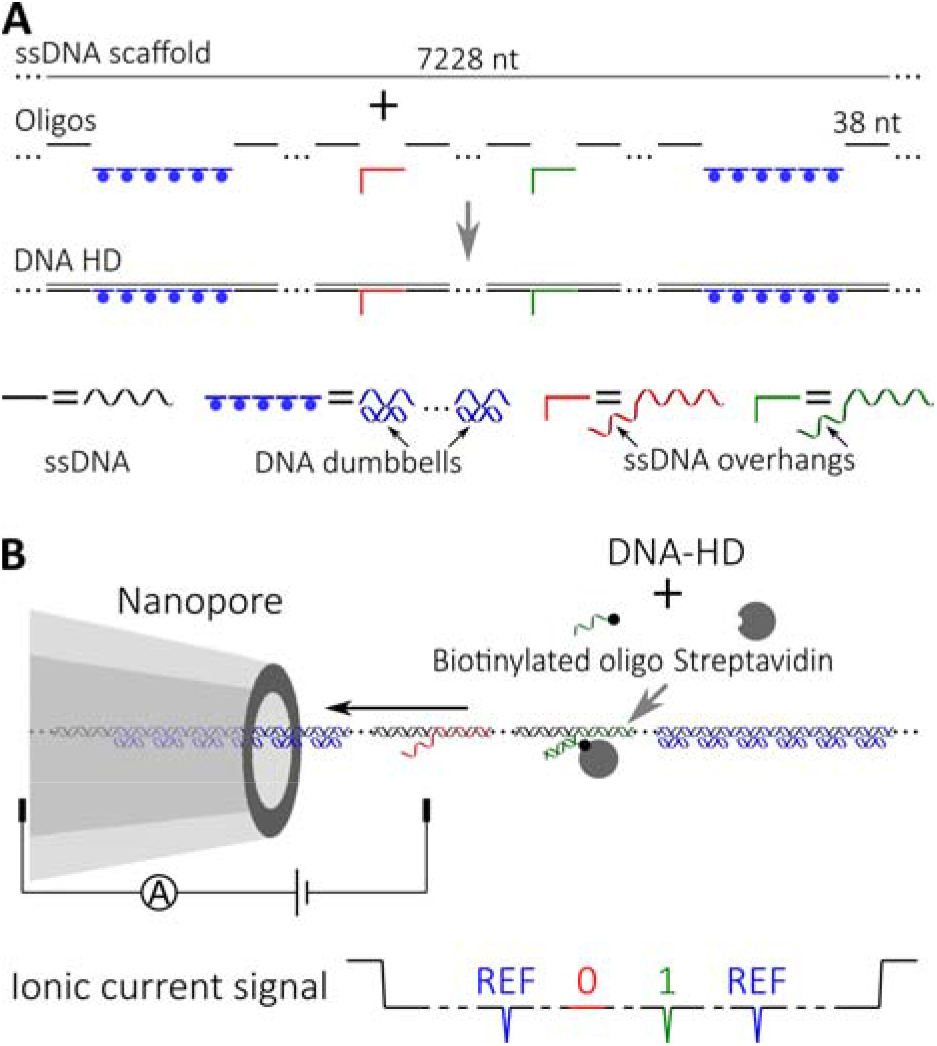
Schematic of the DNA hard drive (DNA-HD). (**A**) Design of the DNA-HD made from ssDNA scaffold and designed short DNA oligonucleotides to form units for data storage and operation. The oligonucleotides assemble to the scaffold via sequence specificity and are designed to incorporate structures like ssDNA overhangs and DNA dumbbells at designed locations. (**B**) Reading information on the DNA-HD using a nanopore. Dumbbells are used as fixed references (REFs) to indicate the storage region. Streptavidin and biotin labelled oligonucleotide (complementary to the ssDNA overhang) is added to specifically bind to the ssDNA overhang to generate detectable signals. Here in the example, only oligonucleotide complementary to green overhang is added, so the green site is read as ‘1’ while the red ‘0’.

Based on this platform, we now demonstrate that we can write, erase and rewrite data on the DNA-HDs. In order to facilitate the proof of concept, the DNA-HD is designed with five sites with ssDNA overhangs between two REF sites (see sequences in Table S2), which we name D1-D5 (Fig.2A). REF sites created by two groups of DNA dumbbells are used to indicate the beginning and end of the data and determine the direction of DNA-HD according to their predesigned positions. Figure 2B shows the schematic workflow of the writing and erasing using D3 as an example. In the initial stage, only a 20 nt overhang partly hybridised with a 10 nt ssDNA blocking strand is on the DNA-HD, which is classified as ‘0’. The 10 nt prevents unintentional writing caused by unavoidable sequence similarity among different sites. Then the information is written by hybridising biotin-labelled ssDNA (Table S3) with the complementary sequence to the overhang (‘WRITE’). Streptavidin is added to bind to biotin to create an observable secondary current peak in nanopore measurements (read as ‘1’). The biotin-labelled strand includes a free toehold end (black) which is used to remove the streptavidin carrying strand from the carrier by adding a complementary strand (Table S3) using a specific strand displacement reaction. This enables the targeted reset of the bit to ‘0’ (‘ERASE’). After this step, the carrier is ready for a new biotin labelled strand to be added to store new information in a new cycle (‘REWRITE’). In the above operations, we waited for 1 hour during each procedure (see details in the Supplementary Text) as a demonstration, which also has potential to be shortened by optimising the experimental parameters (*22, 27, 28*).

**Fig. 2.**
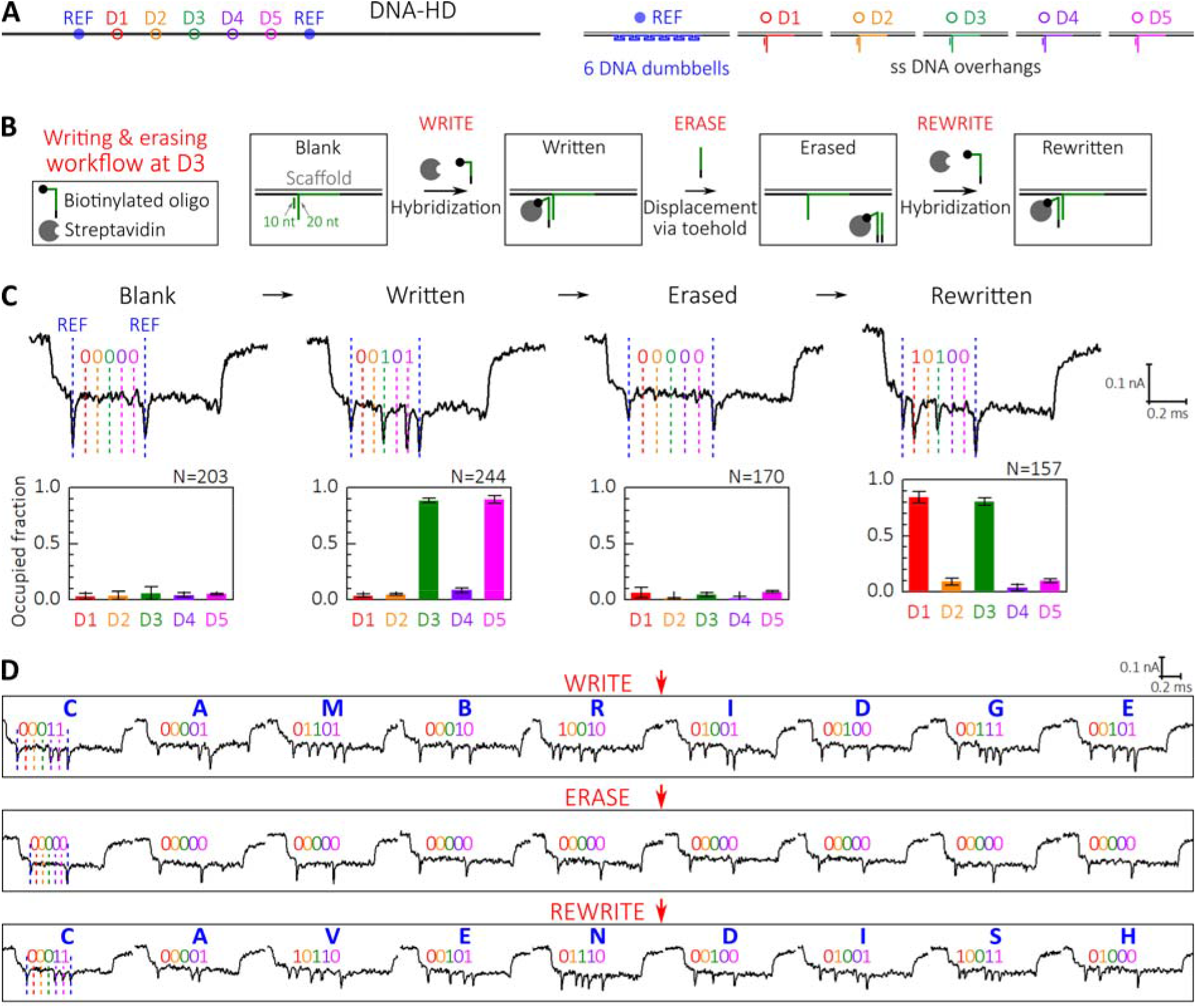
Writing and erasing data on DNA-HDs. (**A**) Design of a writing system on a 7228 bp long DNA-HD. Two groups of DNA dumbbells (marked as “REF” in blue) are used as references to indicate the beginning and end of the storage region. Five positions (D1-D5) with ssDNA overhangs are designated (sequences are shown in Table S2). (**B**) Stages of data writing and erasing on the DNA-HD using D3 as an example. WRITE: A biotin labelled ssDNA strand is used to write as it binds to the overhang with streptavidin added to cause observable signals in the nanopore reading. ERASE: A designed ssDNA strand erases D3 by removing the bound biotin-streptavidin using a strand displacement reaction. REWRITE: After erasing, new data can be encoded by adding new biotinylated ssDNA strand, completing the writing-erasing-rewriting cycle. (**C**) Characterisation of the performance of the system. ‘00101’ was written and erased, and then ‘10100’ was written. Histograms show the statistics from hundreds of molecules (*N* is the number of molecules) where the occupied fraction means the per cent of the signals read as ‘1’ on the site out of the total molecule number. See the statistics in Table S4. (**D**) Demonstration of storing letters as 5-bit ASCII on DNA-HDs and reading the data with nanopores. See the statistics in Table S5.

We characterised the performance of our method by writing ‘00101’ (D1-D5), erasing this information and rewriting ‘10100’. This was done by adding the corresponding oligonucleotides and streptavidin in each step as introduced above and measuring the resulting molecules with nanopores (see the workflow in the Supplementary Text). Figure 2C shows examples of nanopore signals with more events shown in Fig. S1. As each streptavidin label caused a secondary current peak in the event, the information was decoded by analysing the presence or absence of peaks. Our results show a higher signal-to-noise (SNR) ratio compared to a recent paper using MoS_2_ nanopores for the detection of topological variations on DNA (*29*). More importantly, our signals are more consistent and the DNA translocation speed is relatively constant, allowing for correct decoding of the information using single molecules. We analysed hundreds of events and plotted the percentage of detected peaks (named ‘occupied fraction’) at each site (Fig. 2C and Table S4). Initially, all bits had low occupied fractions at the blank state. In the first writing stage, D3 and D5 were written as ‘1’ where the occupied fractions were close to 80% while they were below 10% for other sites. After the information is erased, occupied fractions dropped again for all sites to below 10%. In the second writing stage, occupied fractions for D1 and D3 were also close to 80% and others were below 10%. These significant changes make it possible to correctly detect the information with high certainty using only a few events. It is noticeable that a variation in the translocation velocity can also cause errors if the bits are not correctly assigned, thus placing reasonable REFs and controlling DNA velocity by external parameters like electro-osmotic flow, charge or nanopore geometry will help improve the reading accuracy. Another possibility to reduce errors is to encode ‘0’ with smaller structures (*20*).

After the characterisation, we stored words and changed the data with the rewriting capability (Fig. 2D). Nine samples were prepared and measured. Here we use D1-D5 (corresponding to last 5 bits of ASCII) to encode letters and read them by aggregating the first ten unfolded translocation events for each sample. First, we encoded ‘CAMBRIDGE’ into the samples with example events and codes shown in Figure 3D (details are shown in Fig. S2). Using the same samples, we erased the letters (Fig. S3) and wrote ‘CAVENDISH’ and measured again (Fig. S4). The results show we are able to encode and change data on the DNA-HDs and read the data with nanopores.

**Fig. 3.**
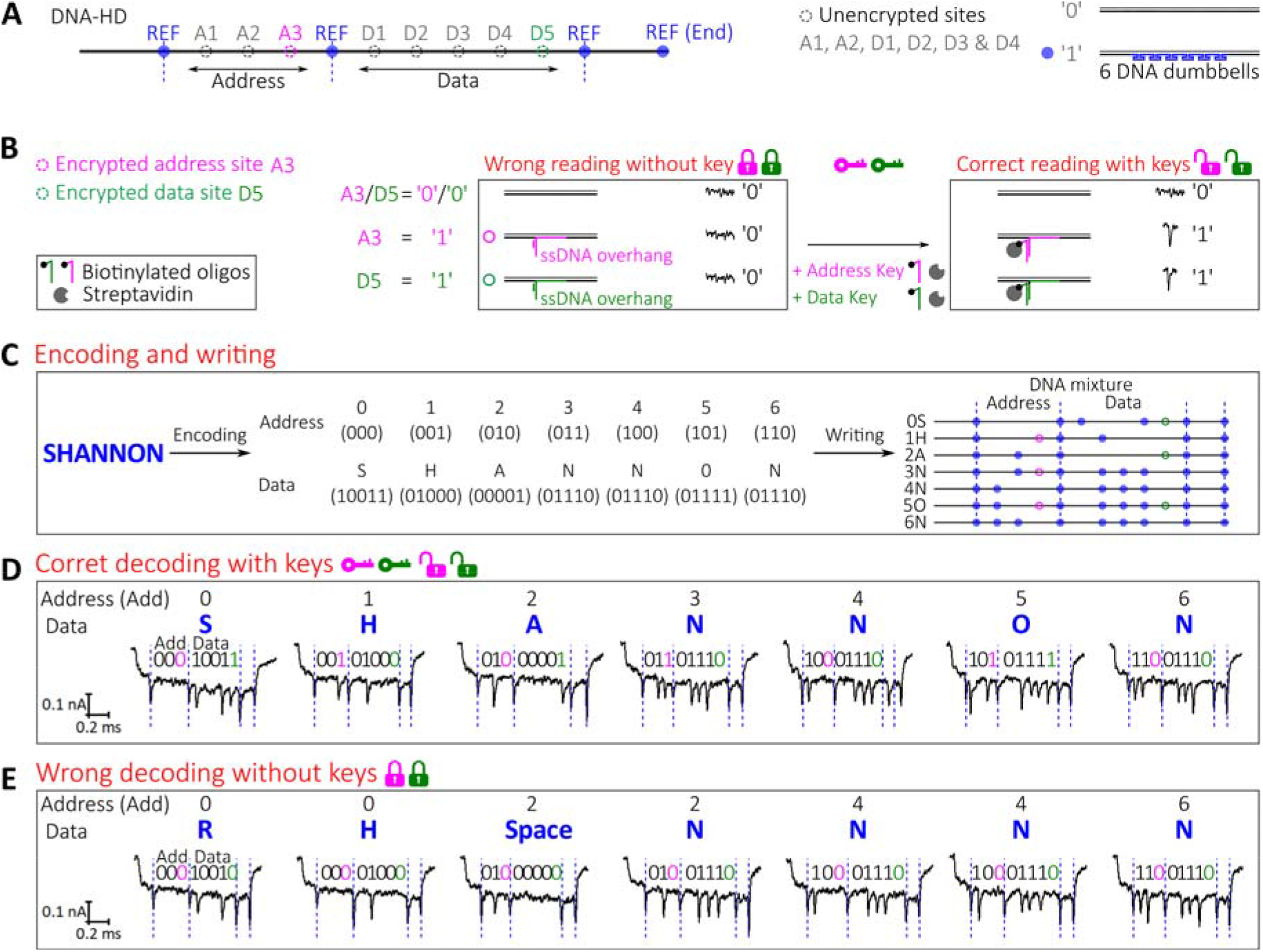
Data encryption on DNA-HDs. (**A**) Design of the DNA-HD for data encryption with 8 bits used for the example of address and data encoding. Schematics of the unencrypted sites are shown, where blank dsDNA is used to represent ‘0’ and dumbbells ‘1’. (**B**) Schematic of the sites for information encryption and the principle of encryption. For these sites, blank dsDNA is used to encode ‘0’ and streptavidin and biotin bound ssDNA overhang to encode ‘1’. But the ‘1’ can only be read with the keys (biotin labelled ssDNA and streptavidin) added. (**C**) An example of data encryption by encoding “SHANNON” into a mixture of DNA-HDs. (**D**) Correct decoding the information with the address and data keys. (**E**) Wrong information decoded without keys. Examples are shown and detailed data analysis is shown in Figures S5 and S6.

Having shown that our DNA-HD can be used as a rewritable storage system, we now encrypt data on ssDNA overhang sites by applying the requirements of additional complementary sequences labelled with biotin and streptavidin for correct reading. The labelled complementary oligonucleotide actually serves as a physical key for data access. Using this idea, we design sites for data encryption on the DNA-HDs (Fig. 3A) where we include the parts for address (A1-A3) and data (D1-D5) split by the REFs (the REF at the end indicates the direction, see sequences in Tables S6 and S7). The address part is used to identify the order of the data when we need to store data in different domains in a mixture. Here we use DNA dumbbells as unencrypted data sites and ssDNA overhangs as encrypted sites (A3 and D5, Fig. 3B), where a blank site (only with ssDNA complementary to the scaffold) means a ‘0’ while a group of dumbbells or the ssDNA overhang a ‘1’. Figure 3B shows the principle of encryption. For these sites, the address key and data key are the streptavidin and biotin labelled oligonucleotides complementary to the overhangs at A3 and D5 respectively. Only when the keys are added can the data be correctly read. An example is shown (Fig. 3C) to encode the word ‘SHANNON’ (to pay tribute to “the father of information theory”) and decode it with nanopores. We encoded each letter in the data part of a DNA-HD and the order in the address, and then made DNA-HDs and mixed them into a mixture. With both address and data keys presented, we successfully decoded the information with example signals shown in Figure 3D (analysis in Fig. S5). However, when the sample was read without the keys, the wrong information appeared (Fig. 3E, details in Fig. S6). First, the order of the letters could not be correctly aligned without the correct address. Second, the encrypted letters on the HDs were not correctly retrieved due to the unrevealed D5. Based on this model, the DNA-HDs can be expanded with more address and data sites on each to encode more practical information. In addition, multiple sites can be chosen as encryption sites to make the data more secure.

In this study, we demonstrate the original innovation only by showing encoding words, but our platform has the full capability to be scaled up using droplet microfluidics which can be automated in parallel to store a large amount of data. Our approach indeed has an inherently lower density compared to data storage in the DNA sequence which is ∼10^6^ of that in hard drives (*2*). In our previous studies, we have demonstrated that our nanopore sensing platform can achieve a resolution of around 100 bp, meaning that we can store one bit per 100 bp and our data density is still ∼3 orders of magnitude higher when compared to hard drives (*20*). Last but not least, our nanostructure basically consists of double-stranded DNA (dsDNA) with nicks, so it should be stable enough, at least more stable than complex DNA nanostructures which can be stored for years (*30-32*), due to the relative long hybridization length of 38 bp for each short strand.

To conclude, we develop an approach to store digital information on addressable molecules along dsDNA, which we name DNA-HD. We can use DNA nanostructures to store data which is fixed when fabricating the molecules, and DNA overhangs flexible for adding or removing molecules as sites for rewritable data storage. Using the ssDNA overhangs, we can encrypt data on the DNA-HDs, and decode by adding the physical keys which are the specific biotin labelled complementary oligonucleotides and streptavidin. Without these keys, one is unable to correctly decode the data. Our method with physical keys and nanopore reading is the most reliable method that can read the information as even sequencing approaches would fail. Storage in the 3D DNA-HDs enables ultra-secure data storage and is promising for applications requiring high security. Compared to the storage in the DNA sequence, our method shows the advantages of easy rewritable capability and secure data storage. Data can be written and operated by adding molecules at room temperature without DNA synthesis. In the future, ever-improving methods under development in DNA nanotechnology will undoubtedly improve both writing capabilities and data security. Secondly, it is also easier to read secondary structures using nanopores than DNA sequencing, and with our nanopore sensing platform it only takes microsecond timescale to read data on one DNA-HD suitable for fast reading. In addition, our system can be used for miniature integration to make an end-to-end DNA storage device (*33*). We assume our method will become a promising route for DNA data storage and possibly also for DNA computing (*34*).

## Acknowledgements

We thank Alexander Ohmann for suggestions on the figures, the lab of Mark Howarth at Oxford University for kindly providing monovalent streptavidin and Boyao Liu from Tsinghua University for some sample preparation.

## Funding

K.C. and U.F.K. acknowledge funding from an ERC Consolidator Grant (Designerpores no. 647144). J.Z. acknowledges funding from an EPSRC grant (EP/M008258/1). F.B. acknowledges funding from Benefactors’ Scholarship and scholars’ research expense scheme from St John’s College, Cambridge.

## Author contributions

K.C., J.Z. and U.F.K. conceived the idea. K.C. and F.B. performed experiments and data analysis. U.F.K supervised the project. K.C. and U.F.K. wrote the manuscript.

## Competing interests

Authors declare no competing interests.

## Data and materials availability

All data needed to evaluate the conclusions in the paper are present in the main text or the supplementary materials.

