## Supplementary Materials for "Secure data storage on DNA hard drives"

**This PDF file includes:**

Materials and Methods
Supplementary Text
Figs. S1 to S6
Tables S1 to S7

**Materials and Methods**

Fabrication of DNA-HD

The DNA HD was made by incubating the linearised M13mp18 single-stranded DNA (ssDNA) scaffold and short 38 bp oligonucleotides with designed positions replaced by those including ssDNA overhangs and DNA dumbbells (purchased from Integrated DNA Technologies with sequences listed Tables S1, S2, S3, S6 and S7).

The 7228 nt scaffold was linearised from M13mp18 ssDNA (7249 bases, N4040S, New England Biolabs) using the following protocol.

1) A 39 nt oligonucleotide (5'-TCTAGAGGATCCCCGGGTACCGAGCTCGAATTCGTAATC-3') was hybridised to the M13mp18 ssDNA by mixing 40 μL M13mp18 ssDNA (250ng/μL), 8 μL 10x Cutsmart buffer (New England Biolabs), 2 μL the 39 nt oligonucleotide (100μM) and 28 μL deionised water.

2) The mixture was heated to 65 °C and linearly cooled down to 25 °C in a thermocycler over 40 minutes.

3) 1 μL of BamHI-HF and 1 μL EcoRI-HF (each 100000 units/ml, New England Biolabs) were added to the reaction mixture followed by incubation at 37 °C for 1 hour.

4) The DNA sample was then immediately purified using a Machery-Nagel NucleoSpin gel and PCR Clean-up kit.

5) The concentration was measured using the NanoDrop and the sample was diluted to a concentration of 100 nM.

The DNA HD was prepared using the following protocol.

1) 8 μL linearised M13mp18 ssDNA (100 nM), 20 μL oligonucleotide mixture (each oligo 200 nM), 4 μL 100 mM $MgCl^2$, 1.2 μL 100 mM Tris-HCl (pH=8), 10 mM EDTA and 6.8 μL deionised water were mixed.

2) The mixture was loaded in a thermocycler and heated to 70°C followed by a linear cooling ramp to 25 °C for 50 minutes.

3) After annealing, these excess oligonucleotides were removed using Amicon Ultra 100 kDa filters. One tube annealed as above was added to 460 μL of 10mM Tris-HCl (pH=8), 0.5 mM $MgCl_2$ and centrifuged at 9000 g for 10 minutes at 4°C. 460 μL more 10mM Tris-HCl (pH=8), 0.5 mM $MgCl_2$ was added and the sample centrifuged again for 10 minutes. The sample was then recovered by turning the filter upside down and centrifuging for 1 minute at 1000 g. This typically yielded ~25 μL at a concentration of ~30-50 ng/uL.

4) Solutions were immediately added after filtering to make the final salt concentration 10 mM Tris-HCl, 100 mM NaCl and 2 mM $MgCl_2$.

Nanopore measurement

Glass nanopores with diameters ~10-15 nm were fabricated by pulling quartz capillaries (outer diameter 0.5 mm and inner diameter 0.2 mm, Sutter Instrument) using a laser-heated pipette puller (P-2000, Sutter Instrument). The fabricated nanopores were assembled into a PDMS chip. DNA sample was diluted in 4 M LiCl , 1×TE (pH=9) with the concentration of 0.2-1 nM and the solution was added to one side of the chip. An Axon Axopatch 200B amplifier (Molecular Devices) was used to apply a voltage 600 mV to drive the DNA through nanopores and measure the ionic current signal. The signal was filtered with an external Bessel filter (Frequency Devices) at 50 kHz and digitized at a 250 kHz sampling rate with a data card (PCI-6251, National Instruments). Data was collected and analysed using home-made LabVIEW algorithms.

**Supplementary Text**

The workflow of the characterisation of the rewritable capability

We used a DNA-HD sample with ssDNA overhangs to start. Oligonucleotides and streptavidin were added to perform the writing and erasing. The resulting molecules were measured with nanopores.

1) Blank.

The DNA concentration of the blank sample was measured as 24 ng/µL (5.04 nM).

2) Writing of '00101'

We mixed the following samples and kept at room temperature for 1 h before nanopore measurement. The ratio of the concentration of the samples is 1 (ssDNA overhang): 4 (biotinylated oligonucleotide):16 (streptavidin).

DNA-HD – blank (5.04 nM): 10 µL

'00101' writing oligonucleotides (containing Oligonucleotides B3 and B5 at 200 nM): 1.01 µL

Monovalent streptavidin (200 nM): 8.08 µL

We name the resulting sample as DNA-HD - '00101' which has a DNA concentration of 2.64 nM.

3) Erasing of '00101'

We mixed the following samples and kept at room temperature for 1 h before nanopore measurement. The ratio of the concentration of the samples is 1 (ssDNA overhang): 8 (erasing oligonucleotide).

DNA-HD - '00101' (2.64 nM): 10 µL

'00101' erasing oligonucleotides (containing Oligonucleotides E3 and E5 at 200 nM): 1.06 µL

We name the resulting sample as DNA-HD – erased which has a DNA concentration of 2.39 nM.

3) Rewriting of '10100'

We mixed the following samples and kept at room temperature for 1 h before nanopore measurement. The ratio of the concentration of the samples is 1 (ssDNA overhang): 13 (erasing oligonucleotide): 52 (streptavidin)

DNA-HD – erased (2.39 nM): 5 µL

'10100' writing oligonucleotides (containing Oligonucleotides B1 and B3 at 200 nM): 0.78 µL

Monovalent streptavidin (200 nM): 6.2 µL

We name the resulting sample as DNA-HD - '10100' which has a DNA concentration of 1.00 nM.

Demonstration of writing 'CAMBRIDGE' – Erasing – rewriting 'CAVENDISH'

The experimental methods are the same as shown above with the corresponding oligonucleotides added in each step and the sample measured with nanopores.

Decoding the data encrypted in DNA-HDs

We prepared the DNA-HD samples encoded with 'S', 'H', 'A', 'N', 'N', 'O' and 'N' with the addresses '0', '1', '2', '3', '4', '5' and '6' respectively. The concentrations are 6.72 nM, 6.51 nM, 5.04 nM, 6.93 nM, 5.88 nM, 7.35 nM and 6.09 respectively. Here we use the ratio of the concentration of the samples 1 (ssDNA overhang): 10 (biotinylated oligonucleotide): 40 (streptavidin) to have high binding efficiency. We mixed the following samples and kept at room temperature for 1 h before nanopore measurement.

1.09 µL, 1.13 µL, 1.46 µL, 1.06 µL, 1.25 µL, 1 µL and 1.21 µL of the seven samples.

2.58 µL 200 nM B5 (sequence shown below), 2.58 µL 200 nM B3 (sequence shown below)

1.5 µL 100 mM $MgCl_2$

4.13 µL 1 µM monovalent streptavidin.

**Fig. S1.**

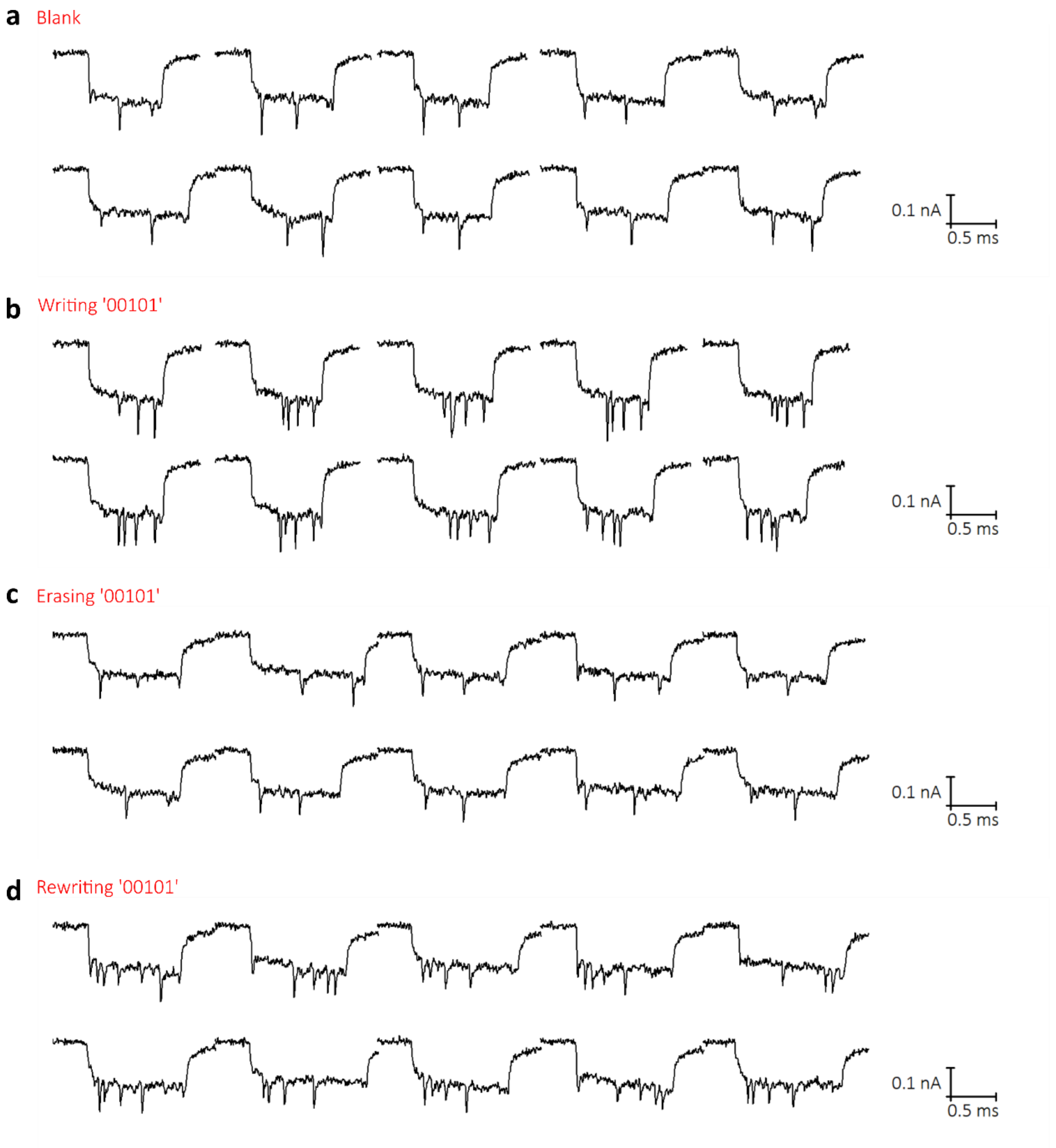


Fig. S1. Example events from nanopore measurement during the writing ‘00101’- erasing – rewriting ‘10100’ process. Events in the four stages are shown in (a)-(d.)

**Fig. S2.**

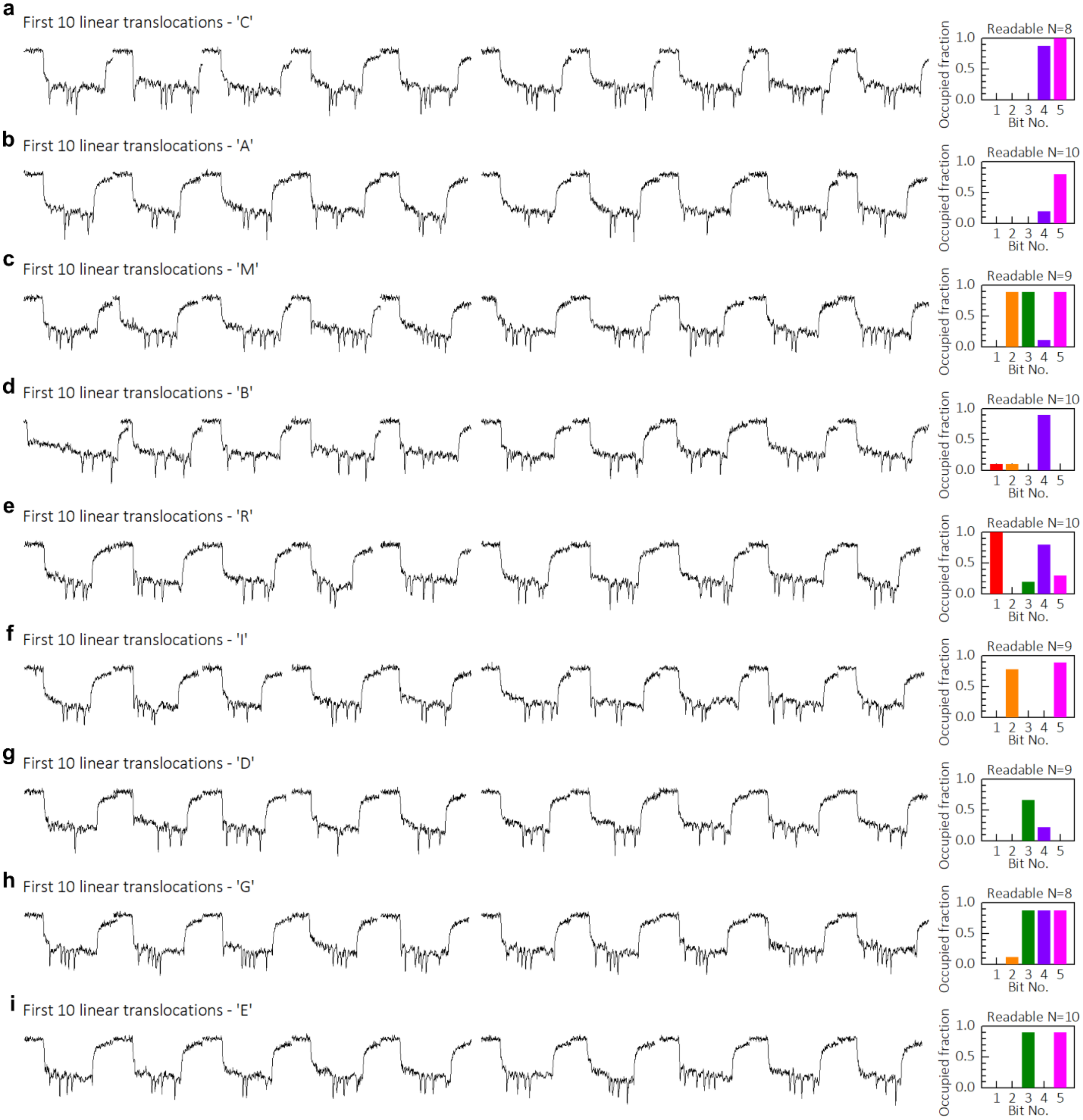


Fig. S2. Nanopore data for the DNA-HDs written with 'CAMBRIDGE'. The first 10 unfolded translocation events and occupied fractions are shown in (a)-(i). In the histogram, we only included the events ('Readable N') with verified correct REF signals.

**Fig. S3.**

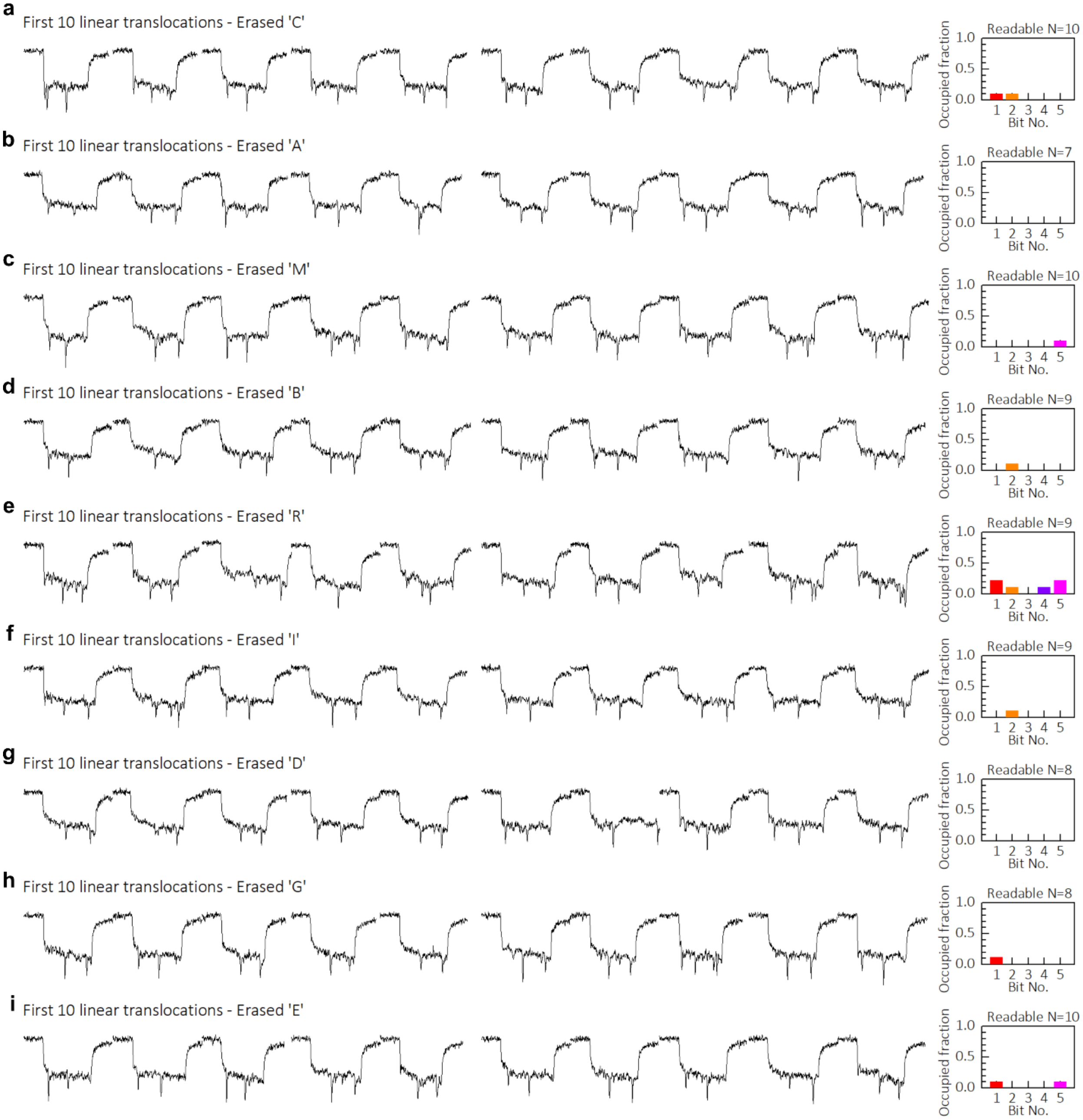


Fig. S3. Nanopore data for the erased DNA-HDs. The first 10 unfolded translocation events and occupied fractions are shown in (a)-(i). In the histogram, we only included the events ('Readable N') with verified correct REF signals.

**Fig. S4.**

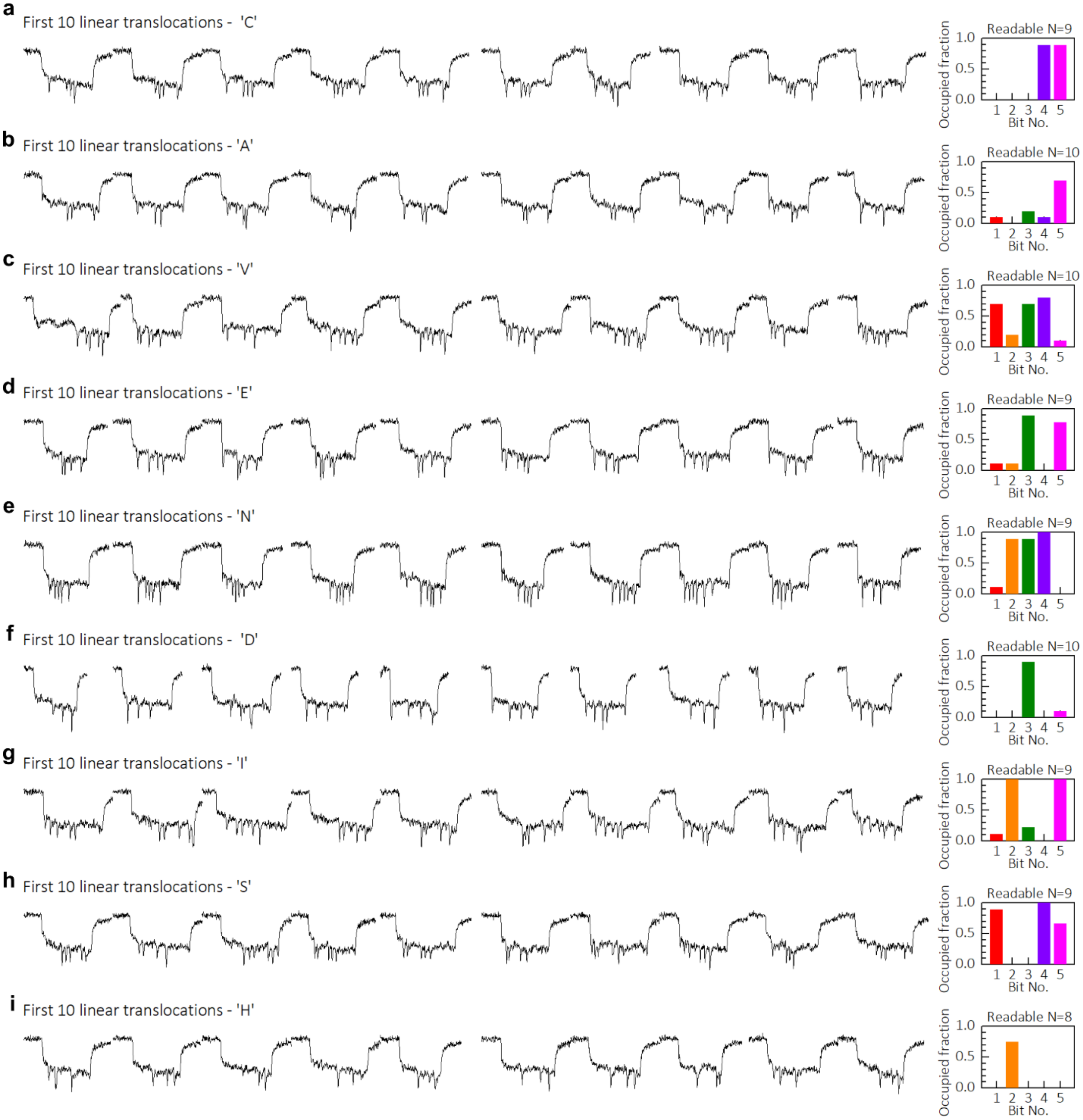


Fig. S4. Nanopore data for the DNA-HDs rewritten with 'CAVENDISH'. The first 10 unfolded translocation events and occupied fractions are shown in (a)-(i). In the histogram, we only included the events ('Readable N') with verified correct REF signals.

**Fig. S5.**

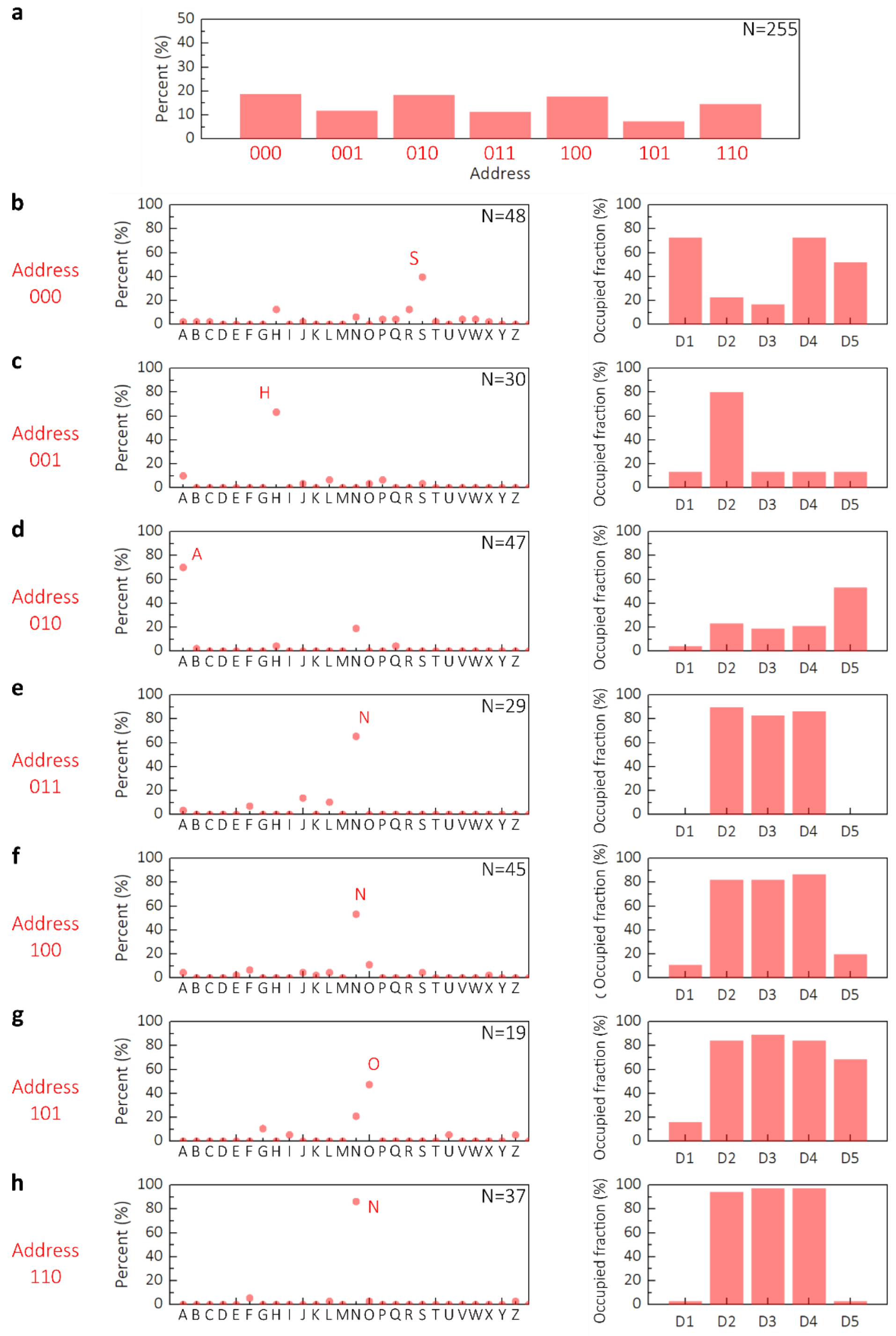


Fig S5. Correct decoding the information with the address and data keys. (a) Per cent of events assigned to addresses 000-100. N is the event number. (b)-(h) show the letter (left) decoded at each address and the occupied fractions at the five data sites.

**Fig. S6.**

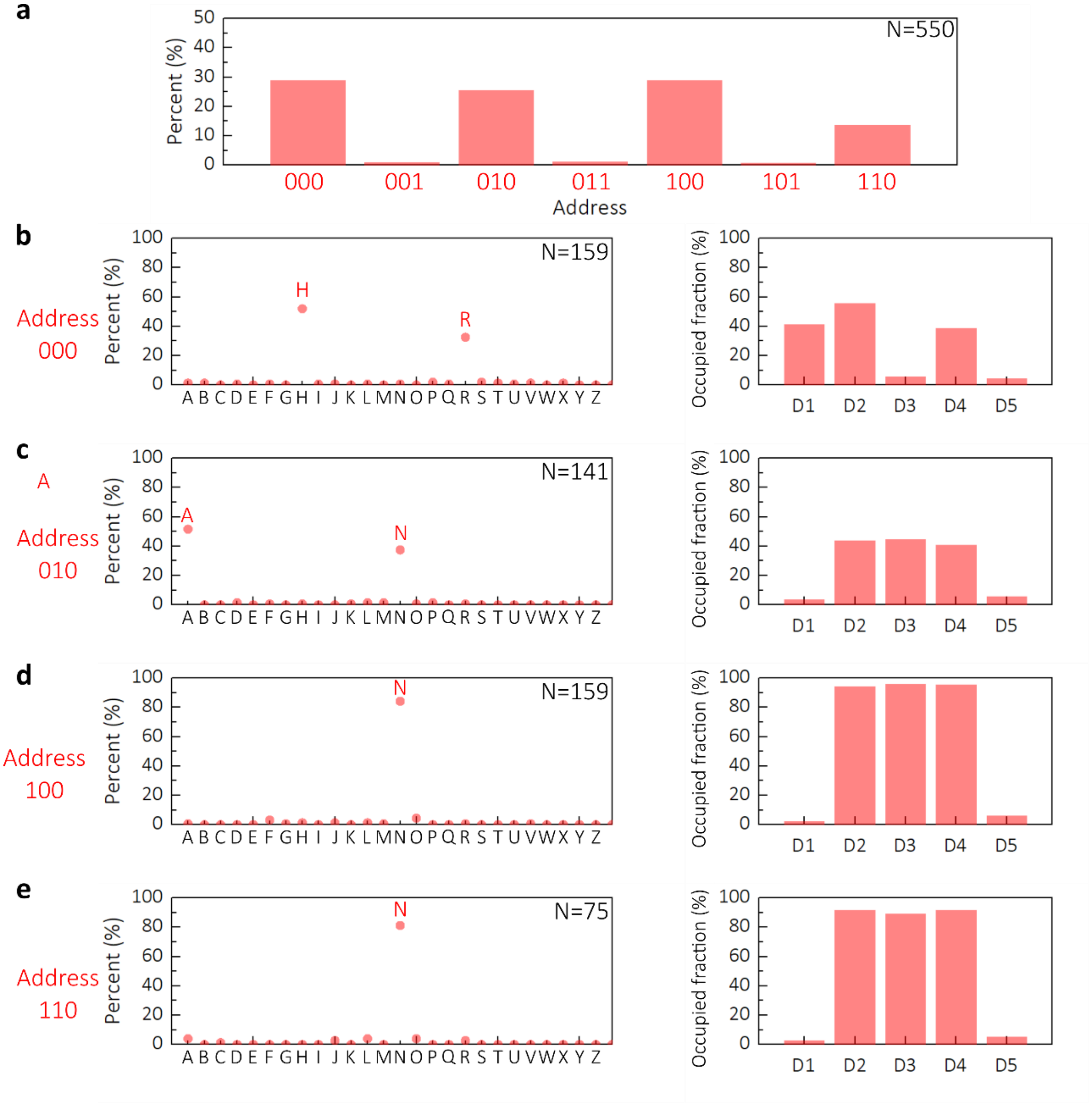


Fig. S6. Wrong information decoded without keys. (a) Per cent of events assigned to addresses 000-100. N is the event number. (b)-(e) show the letter (left) decoded at each address and the occupied fractions at the five data sites. Here we only had four addresses because the third address site was unrevealed so it was always decoded as ‘0’.

### Table S1.

| Oligo No. | Sequence | Oligo No. | Sequence |
|---|---|---|---|
| 1 | TTTTCGTAATCATGGTCATAGCTGTTTCCTGTGTGAAATTGTTATC | 96 | CTTGAGCCATTTGGGAATTAGAGCCAGCAAAATCACCA |
| 2 | CGCTCACAATTCCACACAACATACGAGCCGGAAGCATA | 97 | GTAGCACCATTACCATTAGCAAGGCCGGAAACGTCACC |
| 3 | AAGTGTAAAGCCTGGGGTGCCTAATGAGTGAGCTAACT | 98 | AATGAAACCATCGATAGCAGCACCGTAATCAGTAGCGA |
| 4 | CACATTAATTGCGTTGCGCTCACTGCCCGCTTTCCAGT | 99 | CAGAATCAAGTTTGCCTTTAGCGTCAGACTGTAGCGCG |
| 5 | CGGGAAACCTGTCGTGCCAGCTGCATTAATGAATCGGC | 100 | TTTTCATCGGCATTTTCGGTCATAGCCCCCTTATTAGC |
| 6 | CAACGCGCGGGGAGAGGCGGTTTGCGTATTGGGCGCCA | 101 | GTTTGCCATCTTTTCATAATCAAAATCACCGGAACCAG |
| 7 | GGGTGGTTTTTCTTTTCACCAGTGAGACGGGCAACAGC | 102 | AGCCACCACCGGAACCGCCTCCCTCAGAGCCGCCACCC |
| 8 | TGATTGCCCTTCACCGCCTGGCCCTGAGAGAGTTGCAG | 103 | TCAGAACCGCCACCCTCAGAGCCACCACCCTCAGAGCC |
| 9 | CAAGCGGTCCACGCTGGTTTGCCCCAGCAGGCGAAAAT | 104 | GCCACCAGAACCACCACCAGAGCCGCCGCCAGCATTGA |
| 10 | CCTGTTTGATGGTGGTTCCGAAATCGGCAAAATCCCTT | 105 | CAGGAGGTTGAGGCAGGTCAGACGATTGGCCTTGATAT |
| 11 | ATAAATCAAAAGAATAGCCCGAGATAGGGTTGAGTGTT | 106 | TCACAAACAAATAAATCCTCATTAAAGCCAGAATGGAA |
| 12 | GTTCCAGTTTGGAACAAGAGTCCACTATTAAAGAACGT | 107 | AGCGCAGTCTCTGAATTTACCGTTCCAGTAAGCGTCAT |
| 13 | GGACTCCAACGTCAAAGGGCGAAAAACCGTCTATCAGG | 108 | ACATGGCTTTTGATGATACAGGAGTGTACTGGTAATAA |
| 14 | GCGATGGCCCACTACGTGAACCATCACCCAAATCAAGT | 109 | GTTTTAACGGGGTCAGTGCCTTGAGTAACAGTGCCCGT |
| 15 | TTTTTGGGGTCGAGGTGCCGTAAAGCACTAAATCGGAA | 110 | ATAAACAGTTAATGCCCCCTGCCTATTTCGGAACCTAT |
| 16 | CCCTAAAGGGAGCCCCCGATTTAGAGCTTGACGGGGAA | 111 | TATTCTGAAACATGAAAGTATTAAGAGGCTGAGACTCC |
| 17 | AGCCGGCGAACGTGGCGAGAAAGGAAGGGAAGAAAGCG | 112 | TCAAGAGAAGGATTAGGATTAGCGGGGTTTTGCTCAGT |
| 18 | AAAGGAGCGGGCGCTAGGGCGCTGGCAAGTGTAGCGGT | 113 | ACCAGGCGGATAAGTGCCGTCGAGAGGGTTGATATAAG |
| 19 | CACGCTGCGCGTAACCACCACACCCGCCGCGCTTAATG | 114 | TATAGCCCGGAATAGGTGTATCACCGTACTCAGGAGGT |
| 20 | CGCCGCTACAGGGCGCGTACTATGGTTGCTTTGACGAG | 115 | TTAGTACCGCCACCCTCAGAACCGCCACCCTCAGAACC |
| 21 | CACGTATAACGTGCTTTCCTCGTTAGAATCAGAGCGGG | 116 | GCCACCCTCAGAGCCACCACCCTCATTTTCAGGGATAG |
| 22 | AGCTAAACAGGAGGCCGATTAAAGGGATTTTAGACAGG | 117 | CAAGCCCAATAGGAACCCATGTACCGTAACACTGAGTT |
| 23 | AACGGTACGCCAGAATCCTGAGAAGTGTTTTTATAATC | 118 | TCGTCACCAGTACAAACTACAACGCCTGTAGCATTCCA |
| 24 | AGTGAGGCCACCGAGTAAAAGAGTCTGTCCATCACGCA | 119 | CAGACAGCCCTCATAGTTAGCGTAACGATCTAAAGTTT |
| 25 | AATTAACCGTTGTAGCAATACTTCTTTGATTAGTAATA | 120 | TGTCGTCTTTCCAGACGTTAGTAAATGAATTTTCTGTA |
| 26 | ACATCACTTGCCTGAGTAGAAGAACTCAAACTATCGGC | 121 | TGGGATTTTGCTAAACAACTTTCAACAGTTTCAGCGGA |
| 27 | CTTGCTGGTAATATCCAGAACAATATTACCGCCAGCCA | 122 | GTGAGAATAGAAAGGAACAACTAAAGGAATTGCGAATA |
| 28 | TTGCAACAGGAAAAACGCTCATGGAAATACCTACATTT | 123 | ATAATTTTTTCACGTTGAAAATCTCCAAAAAAAAGGCT |
| 29 | TGACGCTCAATCGTCTGAAATGGATTATTTACATTGGC | 124 | CCAAAAGGAGCCTTTAATTGTATCGGTTTATCAGCTTG |
| 30 | AGATTCACCAGTCACACGACCAGTAATAAAAGGGACAT | 125 | CTTTCGAGGTGAATTTCTTAAACAGCTTGATACCGATA |
| 31 | TCTGGCCAACAGAGATAGAACCCTTCTGACCTGAAAGC | 126 | GTTGCGCCGACAATGACAACAACCATCGCCCACGCATA |
| 32 | GTAAGAATACGTGGCACAGACAATATTTTTGAATGGCT | 127 | ACCGATATATTCGGTCGCTGAGGCTTGCAGGGAGTTAA |
| 33 | ATTAGTCTTTAATGCGCGAACTGATAGCCCTAAAACAT | 128 | AGGCCGCTTTTGCGGGATCGTCACCCTCAGCAGCGAAA |
| 34 | CGCCATTAAAAATACCGAACGAACCACCAGCAGAAGAT | 129 | GACAGCATCGGAACGAGGGTAGCAACGGCTACAGAGGC |
| 35 | AAAACAGAGGTGAGGCGGTCAGTATTAACACCGCCTGC | 130 | TTTGAGGACTAAAGACTTTTTCATGAGGAAGTTTCCAT |
| 36 | AACAGTGCCACGCTGAGAGCCAGCAGCAAATGAAAAAT | 131 | TAAACGGGTAAAATACGTAATGCCACTACGAAGGCACC |
| 37 | CTAAAGCATCACCTTGCTGAACCTCAAATATCAAACCC | 132 | AACCTAAAACGAAAGAGGCAAAAGAATACACTAAAACA |
| 38 | TCAATCAATATCTGGTCAGTTGGCAAATCAACAGTTGA | 133 | CTCATCTTTGACCCCCAGCGATTATACCAAGCGCGAAA |
| 39 | AAGGAATTGAGGAAGGTTATCTAAAATATCTTTAGGAG | 134 | CAAAGTACAACGGAGATTTGTATCATCGCCTGATAAAT |
| 40 | CACTAACAACTAATAGATTAGAGCCGTCAATAGATAAT | 135 | TGTGTCGAAATCCGCGACCTGCTCCATGTTACTTAGCC |
| 41 | ACATTTGAGGATTTAGAAGTATTAGACTTTACAAACAA | 136 | GGAACGAGGCGCAGACGGTCAATCATAAGGGAACCGAA |
| 42 | TTCGACAACTCGTATTAAATCCTTTGCCCGAACGTTAT | 137 | CTGACCAACTTTGAAAGAGGACAGATGAACGGTGTACA |
| 43 | TAATTTTAAAAGTTTGAGTAACATTATCATTTTGCGGA | 138 | GACCAGGCGCATAGGCTGGCTGACCTTCATCAAGAGTA |
| 44 | ACAAAGAAACCACCAGAAGGAGCGGAATTATCATCATA | 139 | ATCTTGACAAGAACCGGATATTCATTACCCAAATCAAC |
| 45 | TTCCTGATTATCAGATGATGGCAATTCATCAATATAAT | 140 | GTAACAAAGCTGCTCATTCAGTGAATAAGGCTTGCCCT |
| 46 | CCTGATTGTTTGGATTATACTTCTGAATAATGGAAGGG | 141 | GACGAGAAACACCAGAACGAGTAGTAAATTGGGCTTGA |
| 47 | TTAGAACCTACCATATCAAAATTATTTGCACGTAAAAC | 142 | GATGGTTTAATTTCAACTTTAATCATTGTGAATTACCT |
| 48 | AGAAATAAAGAAATTGCGTAGATTTTCAGGTTTAACGT | 143 | TATGCGATTTTAAGAACTGGCTCATTATACCAGTCAGG |
| 49 | CAGATGAATATACAGTAACAGTACCTTTTACATCGGGA | 144 | ACGTTGGGAAGAAAAATCTACGTTAATAAAACGAACTA |
| 50 | GAAACAATAACGGATTCGCCTGATTGCTTTGAATACCA | 145 | ACGGAACAACATTATTACAGGTAGAAAGATTCATCAGT |
| 51 | AGTTACAAAATCGCGCAGAGGCGAATTATTCATTTCAA | 146 | TGAGATTTAGGAATACCACATTCAACTAATGCAGATAC |
| 52 | TTACCTGAGCAAAAGAAGATGATGAAACAAACATCAAG | 147 | ATAACGCCAAAAGGAATTACGAGGCATAGTAAGAGCAA |
| 53 | AAAACAAAATTAATTACATTTAACAATTTCATTTGAAT | 148 | CACTATCATAACCCTCGTTTACCAGACGACGATAAAAA |
| 54 | TACCTTTTTTAATGGAAACAGTACATAAATCAATATAT | 149 | CCAAAATAGCGAGAGGCTTTTGCAAAAGAAGTTTTGCC |
| 55 | GTGAGTGAATAACCTTGCTTCTGTAAATCGTCGCTATT | 150 | AGAGGGGGTAATAGTAAAATGTTTAGACTGGATAGCGT |
| 56 | AATTAATTTTCCCTTAGAATCCTTGAAAACATAGCGAT | 151 | CCAATACTGCGGAATCGTCATAAATATTCATTGAATCC |
| 57 | AGCTTAGATTAAGACGCTGAGAAGAGTCAATAGTGAAT | 152 | CCCTCAAATGCTTTAAACAGTTCAGAAAACGAGAATGA |
| 58 | TTATCAAAATCATAGGTCTGAGAGACTACCTTTTTAAC | 153 | CCATAAATCAAAAATCAGGTCTTTACCCTGACTATTAT |
| 59 | CTCCGGCTTAGGTTGGGTTATATAACTATATGTAAATG | 154 | AGTCAGAAGCAAAGCGGATTGCATCAAAAAGATTAAGA |
| 60 | CTGATGCAAATCCAATCGCAAGACAAAGAACGCGAGAA | 155 | GGAAGCCCGAAAGACTTCAAATATCGCGTTTTAATTCG |
| 61 | AACTTTTTCAAATATATTTTAGTTAATTTCATCTTCTG | 156 | AGCTTCAAAGCGAACCAGACCGGAAGCAAACTCCAACA |
| 62 | ACCTAAATTTAATGGTTTGAAATACCGACCGTGTGATA | 157 | GGTCAGGATTAGAGAGTACCTTTAATTGCTCCTTTTGA |
| 63 | AATAAGGCGTTAAATAAGAATAAACACCGGAATCATAA | 158 | TAAGAGGTCATTTTTGCGGATGGCTTAGAGCTTAATTG |
| 64 | TTACTAGAAAAAGCCTGTTTAGTATCATATGCGTTATA | 159 | CTGAATATAATGCTGTAGCTCAACATGTTTTAAATATG |
| 65 | CAAATTCTTACCAGTATAAAGCCAACGCTCAACAGTAG | 160 | CAACTAAAGTACGGTGTCTGGAAGTTTCATTCCATATA |
| 66 | GGCTTAATTGAGAATCGCCATATTTAACAACGCCAACA | 161 | ACAGTTGATTCCCAATTCTGCGAACGAGTAGATTTAGT |
| 67 | TGTAATTTAGGCAGAGGCATTTTCGAGCCAGTAATAAG | 162 | TTGACCATTAGATACATTTCGCAAATGGTCAATAACCT |

| | | | |
|---|---|---|---|
| 68 | AGAATATAAAGTACCGACAAAAGGTAAAGTAATTCTGT | 163 | GTTTAGCTATATTTTCATTTGGGGCGCGAGCTGAAAAG |
| 69 | CCAGACGACGACAATAAACAACATGTTCAGCTAATGCA | 164 | GTGGCATCAATTCTACTAATAGTAGTAGCATTAACATC |
| 70 | GAACGCGCCTGTTTATCAACAATAGATAAGTCCTGAAC | 165 | CAATAAATCATACAGGCAAGGCAAAGAATTAGCAAAAT |
| 71 | AAGAAAAATAATATCCCATCCTAATTTACGAGCATGTA | 166 | TAAGCAATAAAGCCTCAGAGCATAAAGCTAAATCGGTT |
| 72 | GAAACCAATCAATAATCGGCTGTCTTTCCTTATCATTC | 167 | GTACCAAAAACATTATGACCCTGTAATACTTTTGCGGG |
| 73 | CAAGAACGGGTATTAAACCAAGTACCGCACTCATCGAG | 168 | AGAAGCCTTTATTTCAACGCAAGGATAAAAATTTTTAG |
| 74 | AACAAGCAAGCCGTTTTTATTTTCATCGTAGGAATCAT | 169 | AACCCTCATATATTTTAAATGCAATGCCTGAGTAATGT |
| 75 | TACCGCGCCCAATAGCAAGCAAATCAGATATAGAAGGC | 170 | GTAGGTAAAGATTCAAAAGGGTGAGAAAGGCCGGAGAC |
| 76 | TTATCCGGTATTCTAAGAACGCGAGGCGTTTTAGCGAA | 171 | AGTCAAATCACCATCAATATGATATTCAACCGTTCTAG |
| 77 | CCTCCCGACTTGCGGGAGGTTTTGAAGCCTTAAATCAA | 172 | CTGATAAATTAATGCCGGAGAGGGTAGCTATTTTTGAG |
| 78 | GATTAGTTGCTATTTTGCACCCAGCTACAATTTTATCC | 173 | AGATCTACAAAGGCTATCAGGTCATTGCCTGAGAGTCT |
| 79 | TGAATCTTACCAACGCTAACGAGCGTCTTTCCAGAGCC | 174 | GGAGCAAACAAGAGAATCGATGAACGGTAATCGTAAAA |
| 80 | TAATTTGCCAGTTACAAAATAAACAGCCATATTATTTA | 175 | CTAGCATGTCAATCATATGTACCCCGGTTGATAATCAG |
| 81 | TCCCAATCCAAATAAGAAACGATTTTTTGTTTAACGTC | 176 | AAAAGCCCCAAAAACAGGAAGATTGTATAAGCAAATAT |
| 82 | AAAAATGAAAATAGCAGCCTTTACAGAGAGAATAACAT | 177 | TTAAATTGTAAACGTTAATATTTTGTTAAAATTCGCAT |
| 83 | AAAAACAGGGAAGCGCATTAGACGGGAGAATTAACTGA | 178 | TAAATTTTTGTTAAATCAGCTCATTTTTTAACCAATAG |
| 84 | ACACCCTGAACAAAGTCAGAGGGTAATTGAGCGCTAAT | 179 | GAACGCCATCAAAAATAATTCGCGTCTGGCCTTCCTGT |
| 85 | ATCAGAGAGATAACCCACAAGAATTGAGTTAAGCCCAA | 180 | AGCCAGCTTTCATCAACATTAAATGTGAGCGAGTAACA |
| 86 | TAATAAGAGCAAGAAACAATGAAATAGCAATAGCTATC | 181 | ACCCGTCGGATTCTCCGTGGGAACAAACGGCGGATTGA |
| 87 | TTACCGAAGCCCTTTTTAAGAAAAGTAAGCAGATAGCC | 182 | CCGTAATGGGATAGGTCACGTTGGTGTAGATGGGCGCA |
| 88 | GAACAAAGTTACCAGAAGGAAACCGAGGAAACGCAATA | 183 | TCGTAACCGTGCATCTGCCAGTTTGAGGGGACGACGAC |
| 89 | ATAACGGAATACCCAAAAGAACTGGCATGATTAAGACT | 184 | AGTATCGGCCTCAGGAAGATCGCACTCCAGCCAGCTTT |
| 90 | CCTTATTACGCAGTATGTTAGCAAACGTAGAAAATACA | 185 | CCGGCACCGCTTCTGGTGCCGGAAACCAGGCAAAGCGC |
| 91 | TACATAAAGGTGGCAACATATAAAAGAAACGCAAAGAC | 186 | CATTCGCCATTCAGGCTGCGCAACTGTTGGGAAGGGCG |
| 92 | ACCACGGAATAAGTTTATTTTGTCACAATCAATAGAAA | 187 | ATCGGTGCGGGCCTCTTCGCTATTACGCCAGCTGGCGA |
| 93 | ATTCATATGGTTTACCAGCGCCAAAGACAAAAGGGCGA | 188 | AAGGGGGATGTGCTGCAAGGCGATTAAGTTGGGTAACG |
| 94 | CATTCAACCGATTGAGGGAGGGAAGGTAAATATTGACG | 189 | CCAGGGTTTTCCCAGTCACGACGTTGTAAAACGACGGC |
| 95 | GAAATTATTCATTAAAGGTGAATTATCACCGTCACCGA | 190 | CAGTGCCAAGCTTGCATGCCTGCAGGTCGACTCTAGAGGATCTTTT |

Table S1. Sequences of the 190 staples complementary to the scaffold. The length of each oligonucleotide is 38 nt except for the 46 nt ends.

**Table S2.**

| Site | Sequence | To replace Oligo Nos |
|---|---|---|
| REF1 | ACATCACTTGCCTGAGTAGA | 26-30 |
| | AGAACTCAAATCCTCTTTTGAGGAACAAGTTTTCTTGTCTATCGGCCT | |
| | TGCTGGTAATTCCTCTTTTGAGGAACAAGTTTTCTTGTATCCAGAACA | |
| | ATATTACCGCTCCTCTTTTGAGGAACAAGTTTTCTTGTCAGCCATTGC | |
| | AACAGGAAAATCCTCTTTTGAGGAACAAGTTTTCTTGTACGCTCATGG | |
| | AAATACCTACTCCTCTTTTGAGGAACAAGTTTTCTTGTATTTTGACGC | |
| | TCAATCGTCTTCCTCTTTTGAGGAACAAGTTTTCTTGTGAAATGGATT | |
| | ATTTACATTGGCAGATTCAC | |
| | CAGTCACACGACCAGTAATAAAAGGGACAT | |
| D1 | CTCCATTTCCCTTTCATTCT TT TTCGACAACTCGTATTAAATCCTTTGCCCGAACGTTAT | 42 |
| | AGAATGAAAG | |
| D2 | CTCATATCTTCCTATCCTAC TT GTGAGTGAATAACCTTGCTTCTGTAAATCGTCGCTATT | 55 |
| | GTAGGATAGG | |
| D3 | CAACCATCACATCACCAACA TT AGAATATAAAGTACCGACAAAAGGTAAAGTAATTCTGT | 68 |
| | TGTTGGTGAT | |
| D4 | ACCCAAATCTCTGATCTTAC TT TCCCAATCCAAATAAGAAACGATTTTTTGTTTAACGTC | 81 |
| | GTAAGATCAG | |
| D5 | CTATATACTACCTAATACTC TT CATTCAACCGATTGAGGGAGGGAAGGTAAATATTGACG | 94 |
| | GAGTATTAGG | |
| REF2 | TCACAAACAAATAAATCCTCATTAAAGCCAGAATGGAAAGCGCAGTCTCTGAATTT | 106-112 |
| | ACCGTTCCAGTAAGCGTCAT | |
| | ACATGGCTTTTCCTCTTTTGAGGAACAAGTTTTCTTGTTGATGATACA | |
| | GGAGTGTACTTCCTCTTTTGAGGAACAAGTTTTCTTGTGGTAATAAGT | |
| | TTTAACGGGGTCCTCTTTTGAGGAACAAGTTTTCTTGTTCAGTGCCTT | |
| | GAGTAACAGTTCCTCTTTTGAGGAACAAGTTTTCTTGTGCCCGTATAA | |
| | ACAGTTAATGTCCTCTTTTGAGGAACAAGTTTTCTTGTCCCCCTGCCT | |
| | ATTTCGGAACTCCTCTTTTGAGGAACAAGTTTTCTTGTCTATTATTCT | |
| | GAAACATGAAAGTATTAAGA | |
| | GGCTGAGACTCCTCAAGAGAAGGATTAGGATTAGCGGGGTTTTGCTCAGT | |

Table S2. DNA sequences for the design of the rewritable DNA-HD.

**Table S3.**

| Name | Sequence |
| --- | --- |
| B1 | Biotin-TTTTTTT AGAATGAAAGGGAAATGGAG GAGTGAG |
| B2 | Biotin-TTTTTTT GTAGGATAGGAAGATATGAG GGTATGG |
| B3 | Biotin-TTTTTTT TGTTGGTGATGTGATGGTTG AGGAGTG |
| B4 | Biotin-TTTTTTT GTAAGATCAGAGATTTGGGT GTAAGGT |
| B5 | Biotin-TTTTTTT GAGTATTAGGTAGTATATAG TGTAGTG |
| E1 | CTCACTC CTCCATTTCCCTTTCATTCT |
| E2 | CCATACC CTCATATCTTCCTATCCTAC |
| E3 | CACTCCT CAACCATCACATCACCAACA |
| E4 | ACCTTAC ACCCAAATCTCTGATCTTAC |
| E5 | CACTACA CTATATACTACCTAATACTC |

Table S3. Sequences of the oligonucleotides for wiring and erasing data on the rewritable DNA-HD. B1-B5 are biotinylated oligonucleotides that can bind to the overhangs at D1-D5. E1-E5 are oligonucleotides that can bind to B1-B5 to remove them from the DNA-HD using strand displacement reactions.

**Table S4.**

| No. | Pore name | Stage | Encoded information | Total event No. | Unfolded event No. | Readable event No. | Bit 4 occupied No. | Bit 3 occupied No. | Bit 2 occupied No. | Bit 1 occupied No. | Bit 0 occupied No. |
|---|---|---|---|---|---|---|---|---|---|---|---|
| 1 | W&E_1 | 0 (Blank) | 00000 | 693 | 119 | **111** | **3** | **1** | **1** | **5** | **5** |
| 2 | W&E_2 | 0 (Blank) | 00000 | 280 | 49 | **42** | **0** | **1** | **2** | **1** | **2** |
| 3 | W&E_3 | 0 (Blank) | 00000 | 289 | 57 | **50** | **3** | **4** | **6** | **3** | **3** |
| 4 | W&E_4 | 1 (1st write) | 00101 | 478 | 101 | **96** | **3** | **5** | **85** | **6** | **84** |
| 5 | W&E_5 | 1 (1st write) | 00101 | 440 | 80 | **73** | **4** | **4** | **66** | **7** | **64** |
| 6 | W&E_6 | 1 (1st write) | 00101 | 327 | 80 | **75** | **2** | **3** | **65** | **7** | **70** |
| 7 | W&E_7 | 2 (Erase) | 00000 | 373 | 67 | **61** | **7** | **1** | **2** | **2** | **4** |
| 8 | W&E_8 | 2 (Erase) | 00000 | 354 | 75 | **63** | **2** | **1** | **4** | **0** | **5** |
| 9 | W&E_9 | 2 (Erase) | 00000 | 303 | 49 | **46** | **2** | **0** | **2** | **0** | **3** |
| 10 | W&E_10 | 3 (2nd write) | 10100 | 192 | 41 | **35** | **31** | **2** | **28** | **0** | **4** |
| 11 | W&E_11 | 3 (2nd write) | 10100 | 338 | 72 | **61** | **48** | **7** | **47** | **3** | **6** |
| 12 | W&E_12 | 3 (2nd write) | 10100 | 378 | 69 | **61** | **52** | **6** | **51** | **3** | **5** |

Table S4. Statistics of the measurement for the characterization of the writing and erasing (Blank-‘00101’-Erased-‘10100’).

**Table S5.**

| No. | Pore name | Stage | Encoded information | Unfolded events used | Readable event No. | Bit 4 occupied No. | Bit 3 occupied No. | Bit 2 occupied No. | Bit 1 occupied No. | Bit 0 occupied No. |
|---|---|---|---|---|---|---|---|---|---|---|
| 1 | Letter_11 | 1 (1st write) | C (00011) | 10 | **8** | **0** | **0** | **0** | **7** | **8** |
| 2 | Letter_12 | 1 (1st write) | A (00001) | 10 | **10** | **0** | **0** | **0** | **2** | **8** |
| 3 | Letter_13 | 1 (1st write) | M (01101) | 10 | **9** | **0** | **8** | **8** | **1** | **8** |
| 4 | Letter_14 | 1 (1st write) | B (00010) | 10 | **10** | **1** | **1** | **0** | **9** | **0** |
| 5 | Letter_15 | 1 (1st write) | R (10010) | 10 | **10** | **10** | **0** | **2** | **8** | **3** |
| 6 | Letter_16 | 1 (1st write) | I (01001) | 10 | **9** | **0** | **7** | **0** | **0** | **8** |
| 7 | Letter_17 | 1 (1st write) | D (00100) | 10 | **9** | **0** | **0** | **6** | **2** | **0** |
| 8 | Letter_18 | 1 (1st write) | G (00111) | 10 | **8** | **0** | **1** | **7** | **7** | **7** |
| 9 | Letter_19 | 1 (1st write) | E (00101) | 10 | **10** | **0** | **0** | **9** | **0** | **9** |
| 10 | Letter_21 | 2 (Erase) | 00000 | 10 | **10** | **1** | **1** | **0** | **0** | **0** |
| 11 | Letter_22 | 2 (Erase) | 00000 | 10 | **7** | **0** | **0** | **0** | **0** | **0** |
| 12 | Letter_23 | 2 (Erase) | 00000 | 10 | **10** | **0** | **0** | **0** | **0** | **1** |
| 13 | Letter_24 | 2 (Erase) | 00000 | 10 | **9** | **0** | **1** | **0** | **0** | **0** |
| 14 | Letter_25 | 2 (Erase) | 00000 | 10 | **9** | **2** | **1** | **0** | **1** | **2** |
| 15 | Letter_26 | 2 (Erase) | 00000 | 10 | **9** | **0** | **1** | **0** | **0** | **0** |
| 16 | Letter_27 | 2 (Erase) | 00000 | 10 | **8** | **0** | **0** | **0** | **0** | **0** |
| 17 | Letter_28 | 2 (Erase) | 00000 | 10 | **8** | **1** | **0** | **0** | **0** | **0** |
| 18 | Letter_29 | 2 (Erase) | 00000 | 10 | **10** | **1** | **0** | **0** | **0** | **1** |
| 19 | Letter_31 | 3 (2nd write) | C (00011) | 10 | **9** | **0** | **0** | **0** | **8** | **8** |
| 20 | Letter_32 | 3 (2nd write) | A (00001) | 10 | **10** | **1** | **0** | **2** | **1** | **7** |
| 21 | Letter_33 | 3 (2nd write) | V (10110) | 10 | **10** | **7** | **2** | **7** | **8** | **1** |
| 22 | Letter_34 | 3 (2nd write) | E (00101) | 10 | **9** | **1** | **1** | **8** | **0** | **7** |
| 23 | Letter_35 | 3 (2nd write) | N (01110) | 10 | **9** | **1** | **8** | **8** | **9** | **0** |
| 24 | Letter_36 | 3 (2nd write) | D (00100) | 10 | **10** | **0** | **0** | **9** | **0** | **1** |
| 25 | Letter_37 | 3 (2nd write) | I (01001) | 10 | **9** | **1** | **9** | **2** | **0** | **9** |
| 26 | Letter_38 | 3 (2nd write) | S (10011) | 10 | **9** | **8** | **0** | **0** | **9** | **6** |
| 27 | Letter_39 | 3 (2nd write) | H (01000) | 10 | **8** | **0** | **6** | **0** | **0** | **0** |

Table S5. Statistics of the measurement for word storage (‘CAMBRIDGE’ – Erased - ‘CAVENDISH’).

**Table S6.**

| Site | Sequence | To replace Oligo Nos |
|---|---|---|
| REF1 | ACATCACTTGTCCTCTTTTGAGGAACAAGTTTTCTTGTCCTGAGTAGA | 26-30 |
| | AGAACTCAAATCCTCTTTTGAGGAACAAGTTTTCTTGTCTATCGGCCT | |
| | TGCTGGTAATTCCTCTTTTGAGGAACAAGTTTTCTTGTATCCAGAACA | |
| | ATATTACCGCTCCTCTTTTGAGGAACAAGTTTTCTTGTCAGCCATTGC | |
| | AACAGGAAAATCCTCTTTTGAGGAACAAGTTTTCTTGTACGCTCATGG | |
| | AAATACCTACTCCTCTTTTGAGGAACAAGTTTTCTTGTATTTTGACGC | |
| | TCAATCGTCTTCCTCTTTTGAGGAACAAGTTTTCTTGTGAAATGGATT | |
| | ATTTACATTGTCCTCTTTTGAGGAACAAGTTTTCTTGTGCAGATTCAC | |
| | CAGTCACACGACCAGTAATAAAAGGGACAT | |
| REF2 | TGAATCTTACCAACGCTAACGAGCGTCTTTCCAGAGCCTAATTTGCCAGT | 79-85 |
| | TACAAAATAAACAGCCATAT | |
| | TATTTATCCCTCCTCTTTTGAGGAACAAGTTTTCTTGTAATCCAAATA | |
| | AGAAACGATTTCCTCTTTTGAGGAACAAGTTTTCTTGTTTTTGTTTAA | |
| | CGTCAAAAATTCCTCTTTTGAGGAACAAGTTTTCTTGTGAAAATAGCA | |
| | GCCTTTACAGTCCTCTTTTGAGGAACAAGTTTTCTTGTAGAGAATAAC | |
| | ATAAAAACAGTCCTCTTTTGAGGAACAAGTTTTCTTGTGGAAGCGCAT | |
| | TAGACGGGAGTCCTCTTTTGAGGAACAAGTTTTCTTGTAATTAACTGA | |
| | ACACCCTGAACAAAGTCAGA | |
| | GGGTAATTGAGCGCTAATATCAGAGAGATAACCCACAAGAATTGAGTTAAGCCCAA | |
| REF3 | ACAGTTGATTCCCAATTCTGCGAACGAGTA | 161-165 |
| | GATTTAGTTTGACCATTAGA | |
| | TACATTTCGCTCCTCTTTTGAGGAACAAGTTTTCTTGTAAATGGTCAA | |
| | TAACCTGTTTTCCTCTTTTGAGGAACAAGTTTTCTTGTAGCTATATTT | |
| | TCATTTGGGGTCCTCTTTTGAGGAACAAGTTTTCTTGTCGCGAGCTGA | |
| | AAAGGTGGCATCCTCTTTTGAGGAACAAGTTTTCTTGTTCAATTCTAC | |
| | TAATAGTAGTTCCTCTTTTGAGGAACAAGTTTTCTTGTAGCATTAACA | |
| | TCCAATAAATTCCTCTTTTGAGGAACAAGTTTTCTTGTCATACAGGCA | |
| | AGGCAAAGAATTAGCAAAAT | |
| REF4 | CATTCGCCATTCAGGCTGCGCAACTGTTGGGAAG | 186-190 |
| | GGCGATCGGTTCCTCTTTTGAGGAACAAGTTTTCTTGTGCGGGCCTCT | |
| | TCGCTATTACTCCTCTTTTGAGGAACAAGTTTTCTTGTGCCAGCTGGC | |
| | GAAAGGGGGATCCTCTTTTGAGGAACAAGTTTTCTTGTTGTGCTGCAA | |
| | GGCGATTAAGTCCTCTTTTGAGGAACAAGTTTTCTTGTTTGGGTAACG | |
| | CCAGGGTTTTTCCTCTTTTGAGGAACAAGTTTTCTTGTCCCAGTCACG | |
| | ACGTTGTAAATCCTCTTTTGAGGAACAAGTTTTCTTGTACGACGGCCA | |
| | GTGCCAAGCTTCCTCTTTTGAGGAACAAGTTTTCTTGTTGCATGCCTG | |
| | CAGGTCGACTTCCTCTTTTGAGGAACAAGTTTTCTTGTCTAGAGGATCTTTT | |

Table S6. Sequences of the oligonucleotides for forming the dumbbells as REFs on the DNA-HD. Each group consists of 6 DNA dumbbells except for REF4 with 8 DNA dumbbells.

#### Table S7.

| Site | Sequences for '1' | | Sequences for '0' |
|---|---|---|---|
| A1 | CACTAACAACTAATTCCTCTTTTGAGGAACAAGTTTTCTTGTAGATTAGAGC | To replace Oligos 40-43 | Oligos 40-43 |
| | CGTCAATAGATCCTCTTTTGAGGAACAAGTTTTCTTGTTAATACATTT | | |
| | GAGGATTTAGTCCTCTTTTGAGGAACAAGTTTTCTTGTAAGTATTAGA | | |
| | CTTTACAAACTCCTCTTTTGAGGAACAAGTTTTCTTGTAATTCGACAA | | |
| | CTCGTATTAATCCTCTTTTGAGGAACAAGTTTTCTTGTATCCTTTGCC | | |
| | CGAACGTTATTCCTCTTTTGAGGAACAAGTTTTCTTGTTAATTTTAAA | | |
| | AGTTTGAGTAACATTATCATTTTGCGGA | | |
| A2 | TTACCTGAGCAAAAGAAGATGATGAAACAAACATCAAGAAAACA | To replace Oligos 52-57 | Oligos 52-57 |
| | AAATTAATTACATTTAACAA | | |
| | TTTCATTTGATCCTCTTTTGAGGAACAAGTTTTCTTGTATTACCTTTT | | |
| | TTAATGGAAATCCTCTTTTGAGGAACAAGTTTTCTTGTCAGTACATAA | | |
| | ATCAATATATTCCTCTTTTGAGGAACAAGTTTTCTTGTGTGAGTGAAT | | |
| | AACCTTGCTTTCCTCTTTTGAGGAACAAGTTTTCTTGTCTGTAAATCG | | |
| | TCGCTATTAATCCTCTTTTGAGGAACAAGTTTTCTTGTTTAATTTTCC | | |
| | CTTAGAATCCTCCTCTTTTGAGGAACAAGTTTTCTTGTTTGAAAACAT | | |
| | AGCGATAGCTTAGATTAAGA | | |
| | CGCTGAGAAGAGTCAATAGTGAAT | | |
| A3 | CTATATACTACCTAATACTCTTCCAGACGACGACAATAAACAACATGTTCAGCTAATGCA | To replace Oligo 69 | Oligo 69 |
| D1 | CATTCAACCGATTGAGGGAGGGAAGGTCCTCTTTTGAGGAACAAGTTTTCTTGTTAAATATTGA | To replace Oligos 94-97 | Oligos 94-97 |
| | CGGAAATTATTCCTCTTTTGAGGAACAAGTTTTCTTGTTCATTAAAGG | | |
| | TGAATTATCATCCTCTTTTGAGGAACAAGTTTTCTTGTCCGTCACCGA | | |
| | CTTGAGCCATTCCTCTTTTGAGGAACAAGTTTTCTTGTTTGGGAATTA | | |
| | GAGCCAGCAATCCTCTTTTGAGGAACAAGTTTTCTTGTAATCACCAGT | | |
| | AGCACCATTATCCTCTTTTGAGGAACAAGTTTTCTTGTCCATTAGCAAGGCCGGAAACGTCACC | | |
| D2 | TCACAAACAAATAAATCCTCATTAAAGCCAGAATGGAAAGCGCAGTCTCTGAATTT | To replace Oligos 106-112 | Oligos 106-112 |
| | ACCGTTCCAGTAAGCGTCAT | | |
| | ACATGGCTTTTCCTCTTTTGAGGAACAAGTTTTCTTGTTGATGATACA | | |
| | GGAGTGTACTTCCTCTTTTGAGGAACAAGTTTTCTTGTGGTAATAAGT | | |
| | TTTAACGGGGTCCTCTTTTGAGGAACAAGTTTTCTTGTTCAGTGCCTT | | |
| | GAGTAACAGTTCCTCTTTTGAGGAACAAGTTTTCTTGTGCCCGTATAA | | |
| | ACAGTTAATGTCCTCTTTTGAGGAACAAGTTTTCTTGTCCCCCTGCCT | | |
| | ATTTCGGAACTCCTCTTTTGAGGAACAAGTTTTCTTGTCTATTATTCT | | |
| | GAAACATGAAAGTATTAAGA | | |
| | GGCTGAGACTCCTCAAGAGAAGGATTAGGATTAGCGGGGTTTTGCTCAGT | | |
| D3 | TGGGATTTTGCTAAACAACTTT | To replace Oligos 121-124 | Oligos 121-124 |
| | CAACAGTTTCTCCTCTTTTGAGGAACAAGTTTTCTTGTAGCGGAGTGA | | |
| | GAATAGAAAGTCCTCTTTTGAGGAACAAGTTTTCTTGTGAACAACTAA | | |
| | AGGAATTGCGTCCTCTTTTGAGGAACAAGTTTTCTTGTAATAATAATT | | |
| | TTTTCACGTTTCCTCTTTTGAGGAACAAGTTTTCTTGTGAAAATCTCC | | |
| | AAAAAAAAGGTCCTCTTTTGAGGAACAAGTTTTCTTGTCTCCAAAAGG | | |
| | AGCCTTTAATTCCTCTTTTGAGGAACAAGTTTTCTTGTTGTATCGGTTTATCAGCTTG | | |
| D4 | CAAAGTACAACGGAGATTTGTATC | To replace Oligos 134-139 | Oligos 134-139 |
| | ATCGCCTGATAAATTGTGTC | | |
| | GAAATCCGCGTCCTCTTTTGAGGAACAAGTTTTCTTGTACCTGCTCCA | | |
| | TGTTACTTAGTCCTCTTTTGAGGAACAAGTTTTCTTGTCCGGAACGAG | | |
| | GCGCAGACGGTCCTCTTTTGAGGAACAAGTTTTCTTGTTCAATCATAA | | |
| | GGGAACCGAATCCTCTTTTGAGGAACAAGTTTTCTTGTCTGACCAACT | | |
| | TTGAAAGAGGTCCTCTTTTGAGGAACAAGTTTTCTTGTACAGATGAAC | | |
| | GGTGTACAGATCCTCTTTTGAGGAACAAGTTTTCTTGTCCAGGCGCAT | | |
| | AGGCTGGCTGACCTTCATCA | | |
| | AGAGTAATCTTGACAAGAACCGGATATTCATTACCCAAATCAAC | | |
| D5 | CAACCATCACATCACCAACATTAGAGGGGGTAATAGTAAAATGTTTAGACTGGATAGCGT | To replace Oligos 150 | Oligo 150 |

Table S7. Sequences of the oligonucleotides for forming the address and data sites on the DNA-HD.
